# Who rests with whom? Sex composition and group demography shape resting associations in free-ranging dogs

**DOI:** 10.64898/2026.08.28.745223

**Authors:** Sourabh Biswas, Kalyan Ghosh, Kaushikee Sarkar, Anindita Bhadra

**Affiliations:** Behaviour and Ecology Lab, Department of Biological Sciences, Indian Institute of Science Education and Research Kolkata, India; Department of Zoology, University of Gour Banga, India; Department of Environmental Science, University of Calcutta, India; Centre for Ecological Sciences, Indian Institute of Science, Bengaluru, India

**Keywords:** free-ranging dogs, dyadic association, resting behaviour, group composition, social networks, sex-specific sociality

## Abstract

Free-ranging dogs frequently rest near conspecifics, but the demographic factors structuring their resting associations remain poorly understood. We quantified dyadic resting associations in 26 free-ranging dog groups in West Bengal, India, observed between 2019 and 2023. Association strength was estimated from scan-based resting co-occurrences using the Half-Weight Index. We tested whether dyadic association strength varied with dyad sex composition, dyad life-stage composition, group size, and group sex ratio using a generalised additive model for location, scale and shape that accounted for group identity and repeated occurrence of individuals across dyads. Male–male dyads had lower association strengths than female–female dyads, whereas mixed-sex dyads did not differ from female–female dyads. Association strength decreased with increasing group size but increased as the male-to-female ratio within the group increased, while life-stage composition had no detectable effect. Individual-level network metrics, including strength, reach, clustering coefficient, affinity, and eigenvector centrality, did not vary with sex or season. Mixed-sex pairs were also frequently represented among the strongest dyadic associations within groups. These findings indicate that resting associations in free-ranging dogs vary with dyad sex composition and group demography. Further opportunity-controlled analyses are required to determine whether the prominence of mixed-sex dyads reflects preferential association rather than group composition alone.

## Introduction

Free-ranging dog populations are widespread across urban and rural landscapes and represent a dynamic interface between human and ecological systems. These populations show substantial variation in group size, composition, and social behaviour, and their interactions with wildlife and human communities can generate ecological and social challenges (Range & Marshall-Pescini, 2022; Vanak et al., 2009). Understanding how dogs organise socially in such environments is therefore important for behavioural ecology and for informing management in rapidly changing human-dominated landscapes (Biswas, Bhowmik, et al., 2024).

Social behaviour shapes survival and reproduction by influencing access to resources, exposure to risk, and the formation of differentiated relationships within groups. In canids, social organisation varies widely across species and ecological contexts, reflecting differences in resource distribution, competition, and reproductive strategies (Creel et al., 2004; Macdonald, 1983). Domestic dogs, in particular, exhibit flexible social systems across environments, making them a useful model for examining how social structure emerges under variable ecological and demographic conditions. The biological meaning of “association” depends strongly on behavioural context. Proximity during movement or foraging can reflect coordinated activity or shared resource use, whereas proximity during low-activity states may better capture social tole**r**ance and affiliative preferences. Resting behaviour therefore provides a useful context for assessing patterns of association within groups, while also being shaped by ecological constraints such as shelter availability, microclimate, and **l**ocal disturbance (Oliveira et al., 2021; Willis & Brigham, 2007).

Resting association patterns—who rests in close proximity to whom—may reflect stable social preferences and the distribution of tolerance and avoidance within groups. Such associations could be shaped by multiple mechanisms, including kinship (Hamilton, 1964), dominance structure and conflict avoidance (Karavanich & Atema, 1998; Tibbetts et al., 2022), and local ecological conditions that constrain where individuals rest. In free-ranging dogs, however, resting associations have received comparatively less attention than behavioural interactions during activity contexts, and it remains unclear how dyadic association strength is structured within groups and how it varies with group demography and season. m-portantly, a recent population-level field study shows that resting in free-ranging dogs is far from random: dogs show consistent, structured preferences in resting-site selection across seasons, favouring sites near resources with high visibility and relatively low anthropogenic disturbance (Biswas, Ghosh, et al., 2024). This structured “resting template” is likely to shape opportunities for repeated co-occurrence, but whether dyadic resting associations exhibit additional structure consistent with social tolerance and differentiated relationships remains unknown.

Here, we characterise resting association patterns within free-ranging dog groups and test whether dyadic association strength varies with (i) sex combination, (ii) life-stage combination, and (iii) group-level demography (group size and male:female ratio). We also examine whether these dyadic patterns translate into differences in network structure and individual network positions, and whether network measures show seasonal variation at both group and individual levels. We predicted weaker association strength among male–male dyads than among female–female dyads, and relatively strong associations in mixed-sex dyads.

We also predicted that average dyadic association strength would decrease with increasing group size due to dilution of repeated close proximity across more potential partners, and that male:female ratio could influence association patterns by altering the frequency of potential partners and the intensity of within-group competition. Finally, we expected seasonal shifts in association structure if changes in reproductive activity or environmental conditions alter social spacing during resting.

## Materials and Methods

### Study Area

This study was conducted between 2019 and 2023 in West Bengal, India, to investigate resting associations in free-ranging dogs. We focused on three biologically relevant seasons: pre-mating, mating, and post-mating. A total of 26 dog groups were selected based on regional surveys (Sen Majumder et al., 2014) and repeated field observations. Group sizes ranged from 3 to 15 individuals, and across the study we recorded 131 individually identified dogs. Individuals were identified using natural markings and coat colour patterns, supported by repeated sightings.

Study localities were chosen to represent residential, business, and mixed-use neighbourhoods. Part of the study overlapped with the COVID-19 pandemic, and site selection during that period was constrained by travel restrictions. All sites lie within the Indo-Gangetic plain and experience a tropical wet–dry climate, with most rainfall occurring between mid-July and mid-September.

For the purposes of this study, a “group” was defined as a set of dogs that shared the same territory and were repeatedly observed resting and moving within that spatial area. Group composition was not strictly fixed: in some locations membership remained relatively stable across seasons, whereas in others we observed fluctuations over time. Groups were followed across all three seasons within the same annual cycle, allowing seasonal comparison within the same territorial unit.

### Sampling Protocol

Each day was divided into eight 3-hour time blocks: 0000– 0300 h, 0300–0600 h, 0600–0900 h, 0900–1200 h, 1200–1500 h, 1500–1800 h, 1800–2100 h, and 2100–2400 h for observations. A total of 26 groups (Table S1) were sampled across all eight time blocks: four from Bongaon (23.046857, 88.835797) and two from Sodepur (22.700502, 88.379999) in North 24 Parganas district, 13 from Bardhaman Town (23.246974, 87.850792) in Purba Bardhaman district, six from Gayeshpur (22.968002, 88.500352), and one from the IISER Kolkata campus (22.964221, 88.526529) in Nadia district, West Bengal. This scan-based sampling design follows a previously published population-level study of resting behaviour and resting-site selection in free-ranging dogs (Biswas et al., 2024), ensuring methodological comparability. However, whereas that study focused on the physical attributes and placement of resting sites, the present study uses cluster co-occurrence to quantify dyadic resting associations and construct association networks.

Night-time observations (0000–0600 h and 2100–2400 h) were conducted in well-lit street environments, which allowed reliable detection of resting clusters and identification of individuals using coat patterns and natural markings. Because our primary interest was resting proximity (rather than subtle behavioural interactions), visibility under street lighting was sufficient for recording cluster membership and individual identity. Within each season, observations were repeated six times per group per time block, where feasible. The observer followed a predetermined route through the group’s territory. Upon encountering a resting cluster, the observer recorded the identity, sex, and life stage of all dogs present and the cluster composition at that time.

### Definition of Resting Associations and Sampling Unit

A resting cluster was defined as two or more dogs resting within approximately one dog-body length of one another. We applied the gambit-of-the-group approach, whereby all individuals observed within the same resting cluster were considered associated during that observation. Field observations were conducted within predefined 3-hour time blocks. However, for the calculation of association indices in SOCPROG (Whitehead, 2009), each calendar day was defined as one analytical sampling period. All resting-cluster observations recorded during that day were assigned to the same sampling period, while separate resting clusters were retained as distinct groupings within that period. Therefore, dogs recorded in the same resting cluster were classified as observed together, whereas dogs detected on the same day but in different resting clusters were classified as observed in the same sampling period but not together.

Association matrices were constructed from repeated daily sampling periods collected for each focal group. For seasonal comparisons, association data were organised separately for the pre-mating, mating, and post-mating seasons.

### Association Index

To quantify dyadic resting association strength, we used SOCPROG 2.9 (Whitehead, 2008a, 2009) and calculated the Half-Weight Index (HWI) (Cairns & Schwager, 1987), which is commonly used in animal social network studies and is robust to missed detections (Hoppitt & Farine, 2018). For individuals A and B, the HWI was computed as:

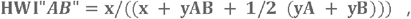

where (x) is the number of daily sampling periods in which (A) and (B) were observed in the same resting cluster, (yAB) is the number of daily sampling periods in which both individuals were observed but in different resting clusters, and (yA) and (yB) are the numbers of daily sampling periods in which only (A) or only (B), respectively, was observed.

### Network Diagnostics and Robustness Checks

To evaluate the reliability and structure of the association data, we examined four diagnostics commonly used in SOCPROG:

#### Correlation coefficient between true and estimated association indices

We assessed the accuracy of estimated associations by calculating the correlation between true and estimated association indices (Whitehead, 2008b). Values close to 1 indicate that estimated indices closely reflect an underlying association structure.

#### Social differentiation

Social differentiation was quantified using the coefficient of variation of true association indices (Whitehead, 2008a), describing how strongly associations vary among dyads (i.e., whether the social structure is relatively homogeneous or strongly differentiated).

#### Cophenetic correlation coefficient

To assess how well hierarchical clustering (dendrograms) represented the association matrix, we calculated the cophenetic correlation coefficient (Bridge, 1993). Higher values indicate a better dendrogram representation of the original association structure.

#### Modularity

We quantified modularity (Newman, 2004) as a measure of whether individuals form distinct clusters with stronger within-cluster associations than expected by chance. Values around or above ~0.3 are often interpreted as suggesting meaningful subdivision.

To assess robustness, we estimated standard errors for correlation and social differentiation using 1000 bootstrap replications. We also implemented Monte Carlo permutation tests to test for preferred or avoided associations (Bejder et al., 1998; Whitehead, 2008b). Permutations were repeated until p-values stabilised, and significance was inferred when the observed association structure (e.g., matrix standard deviation) differed from permuted expectations (p < 0.05).

### Social Structure and Network Metrics

To examine within-group social structure, we identified communities using an eigenvector-based modularity maximization approach (Newman, 2006). Community robustness was evaluated using a community assortativity index (rcom) with 1000 bootstrap iterations, which assesses how consistently individuals are assigned to the same community (Shizuka & Farine, 2016). Networks were visualized in Gephi 0.10.1 (Bastian et al., 2009). When spatial visualization was needed, we used the median latitude/longitude of repeated observations for each individual.

To characterize individual-level positions within networks, we calculated the following weighted network measures:

#### Strength

(Barrat et al., 2004): the sum of an individual’s association indices with all other individuals; higher strength indicates stronger overall connectedness.

#### Eigenvector centrality

(Newman, 2004): measures influence by weighting connections to other well-connected individuals.

#### Reach

(Whitehead, 2008a): quantifies indirect connectedness through third parties, capturing how an individual may be linked to others via intermediate associates.

#### Clustering coefficient

(Holme & Zhao, 2007): measures the extent to which an individual’s associates are also associated with one another in a weighted network.

#### Affinity

Whitehead, 2008a): the strength of an individual’s associates, weighted by association indices, indicating whether an individual tends to associate with well-connected partners. Standard errors for network measures were estimated using 1000 bootstrap replicates

### Statistical Analysis

#### Predictors of dyadic resting association strength

We investigated predictors of dyadic resting association in free-ranging dogs using a generalized additive model for location, scale and shape (GAMLSS) implemented in R (v4.2.0). Dyadic association strength was quantified using the Half-Weight Index (HWI). HWI was modelled as a function of dyad sex combination (female–female, male–female, male– male), dyad life-stage combination (adult–adult, adult– juvenile, adult–pup), group size, and male:female ratio (fixed effects) (Table S1). To account for non-independence in dyadic data, we included random intercepts for group identity and for each individual comprising the dyad (Dog A ID and Dog B ID), thereby controlling for repeated participation of individuals across multiple dyads. Model assumptions and fit were evaluated using standard GAMLSS residual diagnostics (including worm plots and randomized quantile residuals).

#### Sex and seasonal variation in network measures

To test for sex differences in individual network position, we compared five social network measures (strength, reach, clustering coefficient, affinity, and eigenvector centrality) between male and female dogs using Mann–Whitney U tests. A Benjamini–Hochberg correction was applied to control the false discovery rate across the five comparisons.

Seasonal variation in the same network measures, as well as in mean association rate, was examined across three biologically defined seasons based on field observations of reproductive behaviour and local environmental conditions: mating (mid-July to mid-October), post-mating (late October to mid-March), and pre-mating (April to June). For approximately normally distributed variables (strength, affinity, eigenvector centrality, and mean association rate), we used one-way ANOVA; for non-normally distributed variables (reach and clustering coefficient), we used Kruskal–Wallis tests. A Benjamini–Hochberg correction was applied to account for multiple comparisons across network measures. Mean association rate was calculated for each individual as the sum of that individual’s HWI values with all other group members divided by the number of other group members.

#### Sex composition of the strongest dyadic associations

To test whether particular sex combinations were disproportionately represented among the strongest resting associations after accounting for group composition, we identified the single strongest dyad per group (Top-1) and the three strongest dyads per group (Top-3), based on HWI. Each dyad was classified as male–male (MM), male–female (MF), or female– female (FF).

We evaluated deviations from group-specific null expectations using a within-group sex-label permutation procedure. In each permutation, sex labels were randomly reassigned among individuals within the same group while preserving the observed dyads and their HWI values. The Top-1 and Top-3 dyads were then reclassified according to the permuted sex labels, and the numbers of MM, MF, and FF dyads were recorded. This procedure was repeated 5,000 times. For each dyad type, we calculated the mean expected count under the permutation null, the 2.5th and 97.5th percentiles of the null distribution, and a two-sided permutation p-value using a +1 correction to avoid zero p-values.

As a descriptive opportunity-controlled baseline, we also calculated availability-weighted expected counts for each dyad type within each group. The numbers of possible MM, FF, and MF dyads were calculated as m(m−1)/2, f(f−1)/2, and mf, respectively, where m and f represent the numbers of males and females in the group. Group-specific expected counts were then summed across groups for the Top-1 and Top-3 dyad sets. Statistical inference was based on the within-group permutation tests, whereas the availability-weighted expectations were used only as a descriptive comparison.

All analyses were conducted in R.

## Results

### Predictors of dyadic resting association strength

A generalized additive model for location, scale and shape (GAMLSS), including random intercepts for group identity and for the two individuals comprising each dyad, identified significant effects of sex combination, group size, and male:female ratio on dyadic resting association strength (HWI; Table 1). Male–male dyads had significantly lower HWI values than female–female dyads (β = −0.18, SE = 0.07, p < 0.05; Fig. 1), indicating weaker resting associations between males. Male–female dyads did not differ from female– female dyads in their HWI (p = 0.7813; Fig. 1).

**Figure 1.**
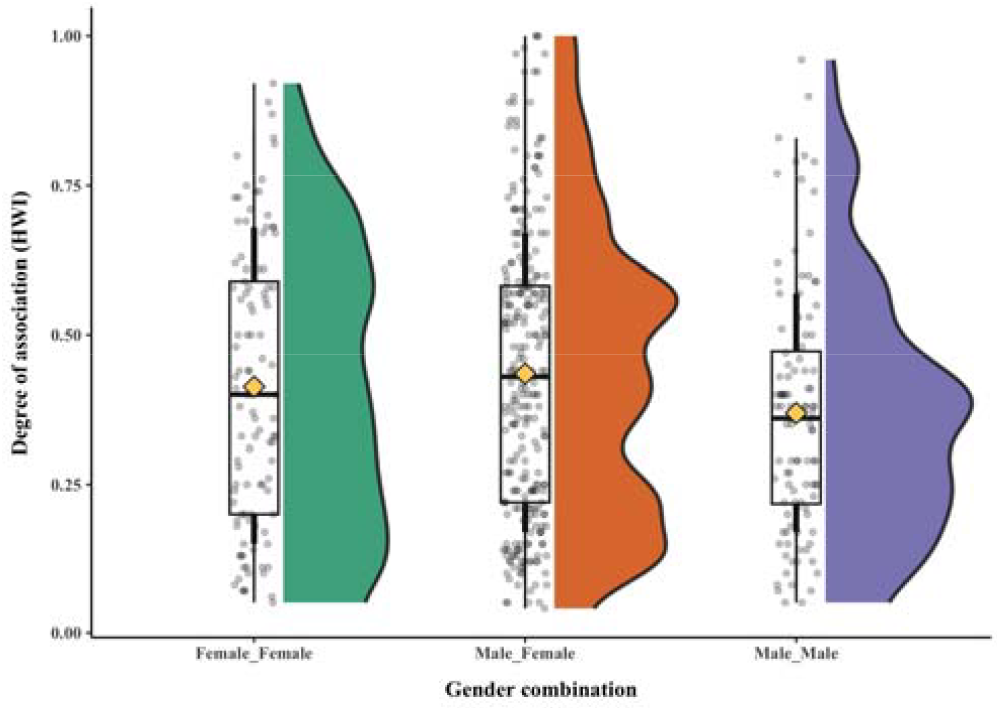
Dyadic resting association strength (HWI) across sex combinations. Shaded half-violins show the distribution of HWI values; boxplots show the median and IQR (whiskers = 1.5×IQR). Points are individual dyads (jittered). Diamonds indicate means.

Group size was strongly negatively associated with HWI (β = −0.06, SE = 0.01, p < 0.001; Fig. 2), such that average dyadic association strength decreased as the number of dogs in a group increased. Male:female ratio had a positive effect on HWI (β = 0.19, SE = 0.07, p < 0.05). Life-stage combination did not significantly predict HWI (adult–juvenile: p = 0.1169; adult–pup: p = 0.1715).

**Figure 2.**
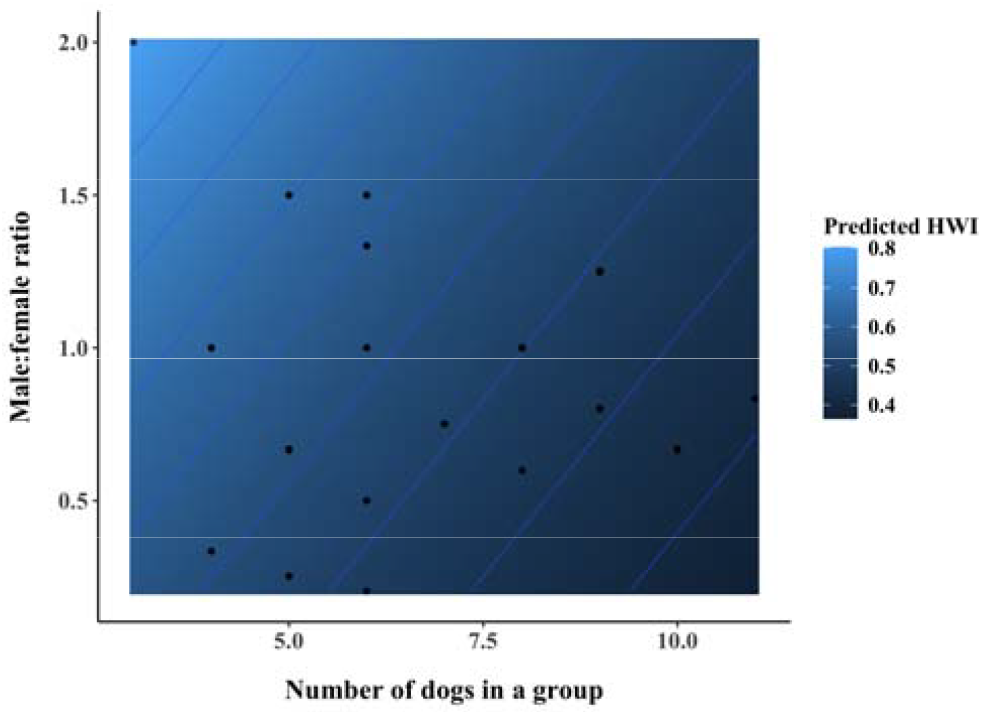
Joint effects of group size and sex ratio on dyadic association strength. The colour surface shows model-predicted Half-Weight Index (HWI) values as a function of the number of dogs in a group and the male:female ratio. Contour lines indicate isopleths of equal predicted HWI. Points show observed group-level combinations of dogs in a group and the male:female ratio used to generate predictions.

### Sex and seasonal variation in network measures

No significant sex differences were detected in ind**iv**idual network measures (Table S2). Strength (Wilcoxon rank-sum: W = 19072, p = 0.5562), reach (W = 18606, p = 0.8736), clustering coefficient (W = 16774, p = 0.2884), and affinit**y (**W = 16769, p = 0.1263) did not differ between males and females. Eigenvector centrality also did not differ significantly between sexes, although it showed a weak, non-significant trend (W = 20354, p = 0.07724). (All comparisons remained non-significant after Benjamini–Hochberg correction.)

The network measures showed no evidence of seasonal variation (Table S3). One-way ANOVAs indicated no differences across seasons in strength (F(2,24) = 0.739, p = 0.488), affinity (F(2,24) = 0.776, p = 0.471), eigenvector centrality (F(2,24) = 2.574, p = 0.0971), or mean association rate (F(2,24) = 0.509, p = 0.607). Kruskal–Wallis tests likewise found no seasonal differences in reach (χ^2^ = 1.1816, df = 2, p = 0.5539) or clustering coefficient (χ^2^ = 0.7314, df = 2, p = 0.6937).

### Sex composition of the strongest dyads

After accounting for group-specific sex composition **u**sing within-group sex-label permutations, the sex composition of the single strongest dyad per group did not differ from null expectations (Table S4; Fig. 3). Among the Top-1 dyads, the observed counts of male–male dyads (MM; observed = 4, null mean = 6.04, 95% null interval = 2–10, p=0.47), male–fe**m**ale dyads (MF; observed = 17, null mean = 14.39, 95% null interval = 9–19, p=0.42), and female–female dyads (FF; observed = 5, null mean = 5.57, 95% null interval = 2–10, p=0.99) were all consistent with their respective null distributions.

**Figure 3.**
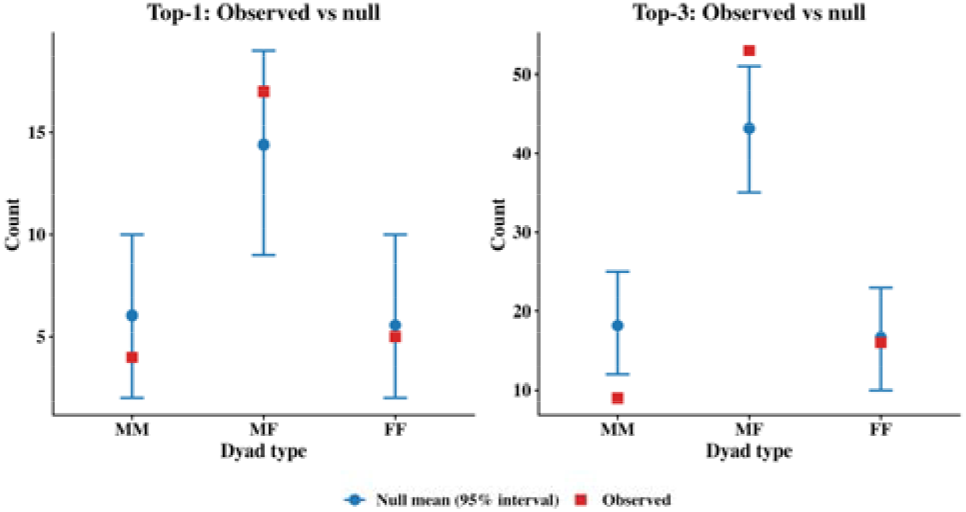
Sex composition of the strongest dyadic relationships. Points indicate the frequency of sex combinations among the single strongest dyad per group (top one) and among the three strongest dyads per group (top three), based on the Half-Weight Index (HWI).

In contrast, the sex composition of the three strongest dyads per group departed from the permutation expectation (Table S4; Fig. 3). MM dyads were under-represented relative to the permutation null (observed = 9, null mean = 18.18, 95% null interval = 12–25, p=0.01), whereas MF dyads were over-represented (observed = 53, null mean = 43.15, 95% null interval = 35–51, p=0.03). FF dyads did not differ from null expectations (observed = 16, null mean = 16.67, 95% null interval = 10–23, p=0.98).

Availability-weighted expected counts closely matched the corresponding permutation null means for both Top-1 and Top-3 dyads (Table S5). These values were used as a descriptive baseline, whereas statistical inference was based on the within-group permutation tests.

## Discussion

The present study shows that resting associations in free-ranging dogs are not random but structured by dyad composition and group demography. Dyadic association strength was low for male–male dyads, as compared to female–female whereas and male–female dyads. Association strength declined strongly with increasing group size and increased with male:female ratio. In contrast, life-stage combination did not predict HWI, and individual-level network measures did not vary significantly between sexes or across seasons. Analyses of the strongest associations further showed that the single strongest dyad per group did not deviate from group-specific permutation expectations, but mixed-sex dyads were overrepresented and male–male dyads were underrepresented among the three strongest dyads within groups. Importantly, resting in free-ranging dogs is known to occur within a structured spatial template—resting-site selection is non-random and consistent across seasons (Biswas et al., 2024)—which likely shapes repeated opportunities for co-occurrence. The results here suggest that, within this ecological template, dyads nonetheless exhibit additional structure associated with sex composition and demography.

The reduced association strength among male–male dyads is consistent with the possibility that male–male resting proximity is constrained by same-sex competition or reduced social tolerance (Clutton-Brock, 1989; Stockley & Bro-Jørgensen, 2011; Trivers, 2017). Similar sex asymmetries have been linked to competitive or aggressive male interactions in other social mammals (Andino et al., 2011; Clutton-Brock & Huchard, 2013). Because these data quantify resting co-occurrence rather than direct behavioural interactions, proximate mechanisms remain uncertain; nevertheless, the pattern is consistent with lower tolerance or higher competitive constraints among males relative to females. Female–female dyads showed stronger associations than male–male dyads, which may reflect greater tolerance and/or more stable partner preferences among females, consistent with broader evidence that females often maintain affiliative relationships that support social cohesion (Silk, 2007). This is also consistent with the observations of allomothering and philopatry in the free-ranging dogs (Paul et al., 2014; Paul & Bhadra, 2018).

Group size had a strong negative effect on dyadic association strength, consistent with a dilution effect: as groups become larger, individuals have more potential partners, so repeated close proximity is distributed across more dyads, reducing average association strength per dyad (Sueur et al., 2011). Group size may also influence spatial spacing and the use of resting locations, reducing repeated co-occurrence among specific pairs (Krause & Ruxton, 2010). In systems where resting sites are selected non-randomly (Biswas et al., 2024), larger groups may additionally be more likely to split across multiple suitable sites, or to redistribute among preferred sites due to crowding or disturbance, further weakening average dyadic co-occurrence. However, because group size also changes the number of possible dyads mathematically, this pattern should be interpreted as reflecting both demographic opportunity and possible behavioural spacing. Together, these results highlight demography as a key determinant of observed association structure.

Male:female ratio also positively predicted HWI, indicating that group sex composition influences resting associations. Sex ratio may alter the opportunity structure for associations, changing the frequency of mixed-sex proximity, or may reflect differences in social stability and spacing behaviour in groups with different compositions. While sexual selection theory predicts that sex ratio can influence social interactions in mixed-sex groups (Andersson, 1994), these data cannot distinguish behavioural mechanisms such as mate guarding, courtship, coalition formation, or aggression. Linking association networks to fine-scale behavioural observations would help clarify why sex ratio affects resting co-occurrence (Garza & Borchert, 1990; Kappel et al., 2017).

Life-stage combination did not significantly predict dyadic association strength, suggesting that resting proximity in this dataset was more strongly associated with sex composition and group demography than with broad age-class combinations. Relatedness and social familiarity could also influence resting patterns (Hamilton, 1964), but kinship was not measured. Similarly, the absence of significant sex differences and seasonal variation in individual-level network measures suggests that males and females occupy broadly similar network positions on average and that network topology is relatively stable across the seasonal periods examined. This does not contradict dyadic sex-composition effects: individuals may have similar overall connectedness while differing in the types of partners that form their strongest associations (Wey et al., 2008; Whitehead, 2008a).

A key insight emerged from analyses of the strongest dyads. When only the single strongest dyad per group was considered, dyad-type composition did not differ from group-specific permutation expectations. However, when the three strongest dyads per group were analysed, mixed-sex dyads were significantly overrepresented and male–male dyads were significantly underrepresented. This suggests that the upper tail of the association-strength distribution contains more mixed-sex associations and fewer male–male associations than expected from group sex composition alone. Importantly, this does not imply that all male–female dyads are stronger than female–female dyads; rather, it suggests that a subset of mixed-sex dyads contributes disproportionately to the strongest resting associations. Such strong mixed-sex associations may reflect reproductive strategies, long-term familiarity, kinship, dominance-related social structure, or other socio-ecological constraints (Hamilton, 1964; Clutton-Brock, 1989; Trivers, 2017; Wittemyer & Getz, 2007; Tibbetts et al., 2022), although these mechanisms cannot be distinguished from co-occurrence data alone.

A limitation of the present approach is that co-occurrence within resting clusters cannot fully separate social preference from repeated convergence on preferred resting sites. Given that resting-site selection is structured and consistent (Biswas et al., 2024), site-driven aggregation may contribute to observed associations, particularly in resource-rich or highly preferred locations. However, the consistently high occurrence of mixed sex dyads in the bootstrapped data and lower than expected occurrence of male-male dyads lends strength to our conclusions. Future work could address this limitation by incorporating resting-site identity as a random effect, modelling dyadic co-occurrence conditional on site use, or combining association data with GPS/proximity loggers and fine-scale behavioural observations.

Overall, resting associations in free-ranging dogs are shaped by sex composition and group demography, with weaker male–male dyadic associations, strong group-size effects, and a disproportionate representation of mixed-sex dyads among the three strongest associations within groups. Resting proximity therefore provides a useful behavioural context for describing social structure in a widespread, human-associated carnivore and for generating testable hypotheses about the behavioural mechanisms underlying differentiated associations.

## Acknowledgements

We thank the Indian Institute of Science Education and Research (IISER) Kolkata for infrastructural support. We also thank Dr Udipta Chakraborty for insightful suggestions during the analysis. This paper was typeset with the bioRxiv word template by @Chrelli: www.github.com/chrelli/bioRxiv-word-template

## Author contributions

S.B., K.G., and K.S. conducted fieldwork. S.B. curated the data, performed the analyses, and drafted the manuscript. S.B. and A.B. conceived the study. A.B. supervised the work, secured funding, and reviewed and edited the manuscript.

## Funding

S.B. was supported by a fellowship from the University Grants Commission (UGC), India. This work was supported by IISER Kolkata.

## Data availability

Data and code supporting the findings of this study will be deposited in a public repository upon acceptance.

## Competing interest statement

The authors declare no competing interests.

## Supplementary materials

**Table S1.** Group-level demographic composition reconstructed from the dyadic association dataset. For each free-ranging dog group, the table reports the number of unique females and males represented in the dyadic dataset, the resulting group size, and the male:female ratio. Group size was calculated as the total number of unique identified females and males within each group, and the male:female ratio was calculated as the number of males divided by the number of females. These reconstructed values were used as a demographic consistency check against the group-level covariates included in the dyadic association analysis.

| Group name | Location | Female | Male | Group size | Male-female ratio |
| --- | --- | --- | --- | --- | --- |
| bng_bsf | Bongaon | 4 | 3 | 7 | 0.75 |
| bng_firebgd | Bongaon | 7 | 8 | 15 | 1.14 |
| bng_netajimk | Bongaon | 5 | 6 | 11 | 1.20 |
| bng_sbi | Bongaon | 4 | 5 | 9 | 1.25 |
| bwn_bedclg | Bardhaman | 5 | 5 | 10 | 1.00 |
| bwn_goda | Bardhaman | 4 | 5 | 9 | 1.25 |
| bwn_goda_shibtala_a | Bardhaman | 2 | 7 | 9 | 3.50 |
| bwn_kajirhat | Bardhaman | 4 | 3 | 7 | 0.75 |
| bwn_ksp_east | Bardhaman | 3 | 3 | 6 | 1.00 |
| bwn_ksp_n | Bardhaman | 1 | 2 | 3 | 2.00 |
| bwn_ksp_west | Bardhaman | 3 | 4 | 7 | 1.33 |
| bwn_med_col | Bardhaman | 4 | 5 | 9 | 1.25 |
| bwn_mithapukur | Bardhaman | 4 | 5 | 9 | 1.25 |
| bwn_mohanbagan | Bardhaman | 6 | 6 | 12 | 1.00 |
| bwn_tarabagh | Bardhaman | 5 | 5 | 10 | 1.00 |
| bwn_tikarhat | Bardhaman | 3 | 3 | 6 | 1.00 |
| bwn_university_gate | Bardhaman | 4 | 1 | 5 | 0.25 |
| gp_kalimandr | Gayeshpur | 4 | 5 | 9 | 1.25 |
| gp_nabarun | Gayeshpur | 4 | 2 | 6 | 0.50 |
| gp_ns_field | Gayeshpur | 5 | 2 | 7 | 0.40 |
| gp_shanti | Gayeshpur | 4 | 4 | 8 | 1.00 |
| gp_townclub | Gayeshpur | 4 | 5 | 9 | 1.25 |
| gp_word1 | Gayeshpur | 3 | 2 | 5 | 0.67 |
| iiserk_rc | IISER Kolkata Campus | 3 | 4 | 7 | 1.33 |
| sd_gorbhobani | Sodepur | 2 | 2 | 4 | 1.00 |
| sd_nilachal | Sodepur | 2 | 4 | 6 | 2.00 |

**Table S2.**
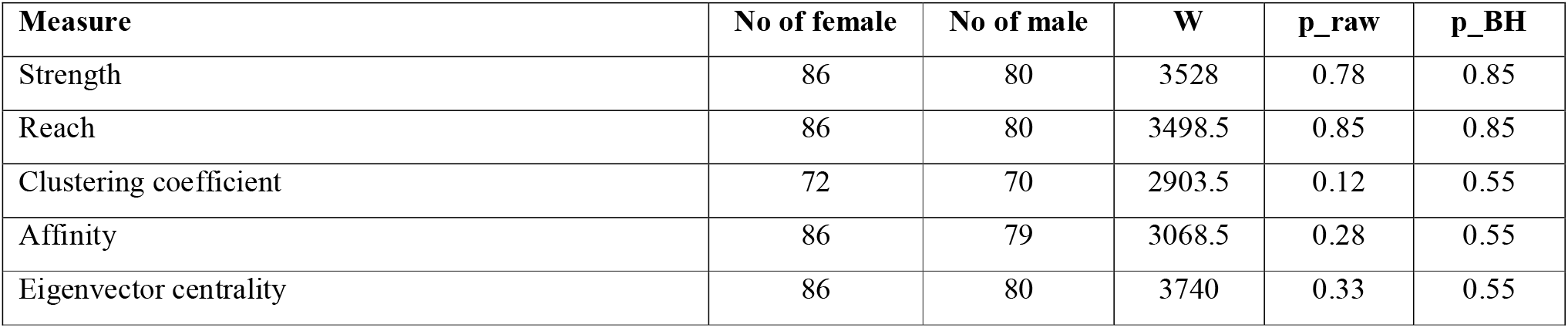
Comparison of individual-level social network metrics between female and male free-ranging dogs. Sex differences in weighted network strength, reach, clustering coefficient, affinity, and eigenvector centrality were tested using Wilcoxon rank-sum tests. Sample sizes for females and males are reported separately for each metric because missing values differed among measures. W denotes the Wilcoxon test statistic, p_raw is the unadjusted p-value, and p_BH is the p-value after Benjamini– Hochberg correction for the five network-metric comparisons.

**Table S3.** Seasonal variation in group-level social network metrics of free-ranging dogs. Seasonal differences among the pre-mating, mating, and post-mating periods were tested using one-way ANOVA for strength, affinity, eigenvector centrality, and mean association rate, and Kruskal–Wallis tests for reach and clustering coefficient. The table reports the test statistic, numerator and denominator degrees of freedom where applicable, unadjusted p-values (p_raw), and Benjamini–Hochberg-adjusted p-values (p_BH) across the network-metric comparisons. Mean association rate was analysed separately and was therefore not included in the multiple-testing correction.

| Measure | Test | Statistic | df1 | df2 | $p_{\text{raw}}$ | $p_{\text{BH}}$ |
| --- | --- | --- | --- | --- | --- | --- |
| Strength | One-way ANOVA | 0.74 | 2 | 24 | 0.49 | 0.69 |
| Affinity | One-way ANOVA | 0.78 | 2 | 24 | 0.47 | 0.69 |
| Eigenvector centrality | One-way ANOVA | 2.57 | 2 | 24 | 0.10 | 0.49 |
| Mean association rate | One-way ANOVA | 0.51 | 2 | 24 | 0.61 | NA |
| Reach | Kruskal Wallis | 1.18 | 2 | NA | 0.55 | 0.69 |
| Clustering coefficient | Kruskal Wallis | 0.73 | 2 | NA | 0.69 | 0.69 |

**Table S4.** Sex composition of the strongest dyadic resting associations compared with within-group permutation expectations. For each group, the single strongest dyad (Top-1) and the three strongest dyads (Top-3), based on the Half-Weight Index, were classified as male–male (MM), male–female (MF), or female–female (FF). Sex labels were permuted among individuals within each group 5,000 times while retaining the observed dyads and HWI values. The table reports the observed number of dyads, the mean expected count under the permutation null, the 2.5th and 97.5th percentiles of the null distribution, and the two-sided permutation p-value.

| Rank set | Dyad type | Observed | Null mean | Null 2.5<sup>th</sup> | Null 97.5<sup>th</sup> | $p_{\text{two-sided}}$ |
| --- | --- | --- | --- | --- | --- | --- |
| Top-1 | MM | 4 | 6.04 | 2 | 10 | 0.47 |
| Top-1 | MF | 17 | 14.39 | 9 | 19 | 0.42 |
| Top-1 | FF | 5 | 5.57 | 2 | 10 | 0.99 |
| Top-3 | MM | 9 | 18.18 | 12 | 25 | 0.01 |
| Top-3 | MF | 53 | 43.15 | 35 | 51 | 0.03 |
| Top-3 | FF | 16 | 16.67 | 10 | 23 | 0.98 |

**Table S5.** Availability-weighted expected counts of dyad sex combinations among the strongest resting associations. Expected numbers of male–male (MM), male–female (MF), and female–female (FF) dyads were calculated separately for the single strongest dyad per group (*K = 1*) and the three strongest dyads per group (*K = 3*). For each group, the expected proportion of each dyad type was based on the number of possible dyads of that type given the observed numbers of males and females; group-specific expectations were then summed across all groups.

| K | Expected MM | Expected MF | Expected FF |
| --- | --- | --- | --- |
| 1 | 6.05 | 14.39 | 5.56 |
| 3 | 18.14 | 43.17 | 16.69 |

## GAMLSS Diagnostic plots

**Figure S1.**
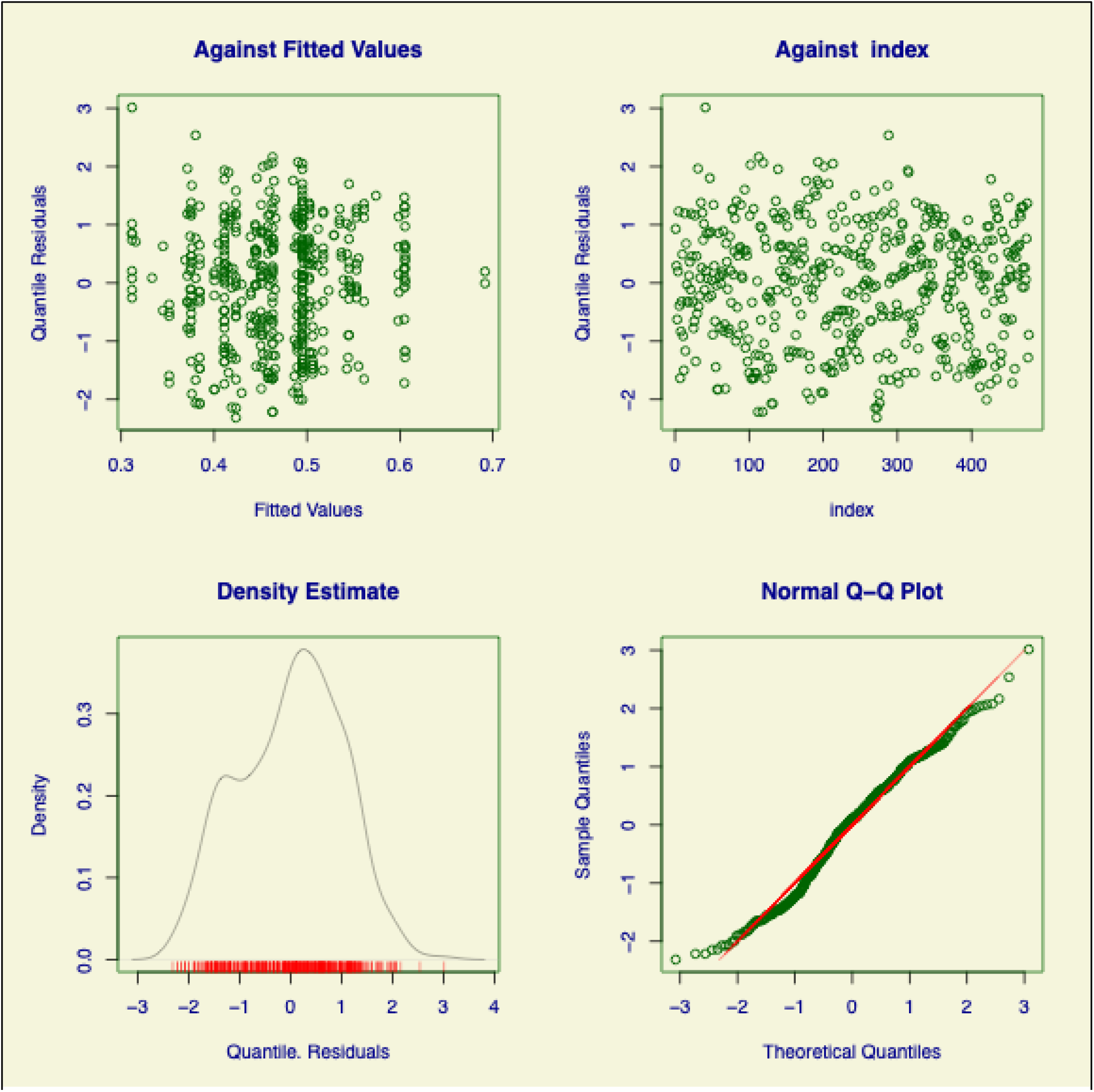
Standard diagnostic plots for the final GAMLSS model of dyadic resting association strength. The panels show randomized quantile residuals plotted against fitted values and observation index, the estimated residual-density distribution, and a normal Q–Q plot. Residuals were broadly centred around zero with no strong systematic trend across fitted values or observation order. The Q–Q plot showed an approximately linear pattern through most of the distribution, with some departures in the tails.

**Figure S2.**
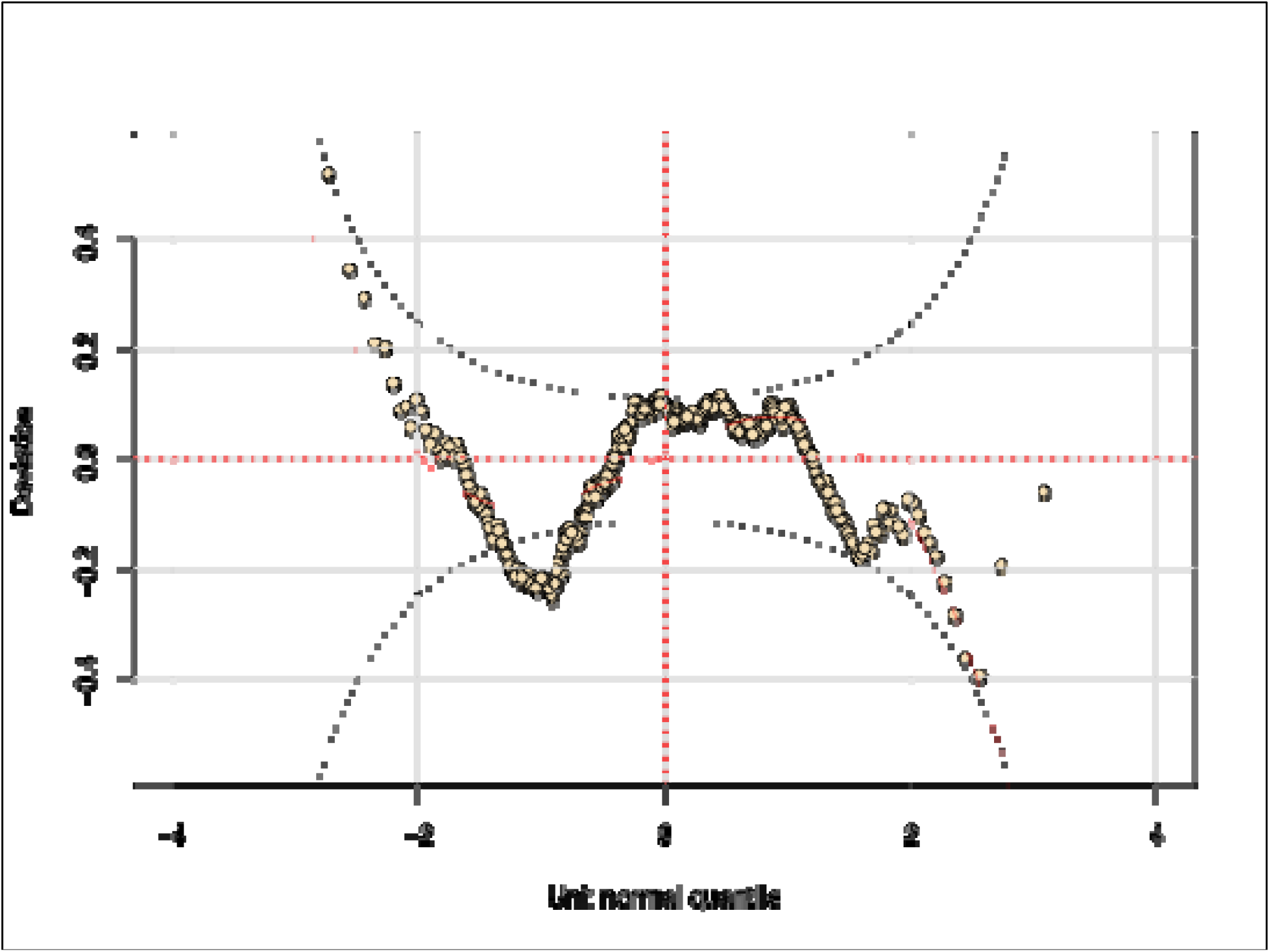
Worm plot for the final GAMLSS model of dyadic resting association strength. The worm plot displays deviations of the randomized quantile residuals from the expected unit-normal distribution across theoretical quantiles. The solid red curve indicates the smoothed residual pattern, the horizontal reference line represents zero deviation, and the dashed curves show the approximate confidence limits. The curved pattern and several points outside the confidence envelope, particularly toward the tails, indicate some remaining lack of fit in the residual distribution.

## Notes

### Competing Interest Statement

The authors have declared no competing interest.

## References

Andersson, M. (1994). Sexual selection (Vol. 72). Princeton University Press.

Andino, N., Reus, L., Cappa, F. M., Campos, V. E., & Giannoni, S. M. (2011). Social environment and agonistic interactions: strategies in a small social mammal. Ethology, 117(11), 992–1002.

Barrat, A., Barthelemy, M., Pastor-Satorras, R., & Vespignani, A. (2004). The architecture of complex weighted networks. Proceedings of the National Academy of Sciences, 101(11), 3747–3752.

Bastian, M., Heymann, S., & Jacomy, M. (2009). Gephi: an open source software for exploring and manipulating networks. Proceedings of the International AAAI Conference on Web and Social Media, 3(1), 361–362.

Bejder, L., Fletcher, D., & Bräger, S. (1998). A method for testing association patterns of social animals. Animal Behaviour, 56(3), 719–725. 10.1006/anbe.1998.0802

Biswas, S., Bhowmik, T., Ghosh, K., Roy, A., Lahiri, A., Sarkar, S., & Bhadra, A. (2024). Scavengers in the human-dominated landscape: an experimental study. Philosophical Transactions of the Royal Society B: Biological Sciences, 379(1909). 10.1098/rstb.2023.0179

Biswas, S., Ghosh, K., Sarkar, K., Benny, L., Katti, M., & Bhadra, A. (2024). A population-level study reveals hidden patterns in resting site choice of free-ranging dogs. Biological Journal of the Linnean Society, 143(3), blae095. 10.1093/biolinnean/blae095

Bridge, P. D. (1993). Classification. In J. C. Fry (Ed.), Biological Data Analysis (pp. 219–242). Oxford University Press.

Cairns, S. J., & Schwager, S. J. (1987). A comparison of association indices. Animal Behaviour, 35(5), 1454–1469.

Clutton-Brock, T. H. (1989). Review lecture: mammalian mating systems. Proceedings of the Royal Society of London. B. Biological Sciences, 236(1285), 339–372.

Clutton-Brock, T. H., & Huchard, E. (2013). Social competition and selection in males and females. Philosophical Transactions of the Royal Society B: Biological Sciences, 368(1631), 20130074.

Creel, S., Mills, M. G. L., & McNutt, J. W. (2004). Demography and population dynamics of African wild dogs in three critical populations. Biology and Conservation of Wild Canids, 12, 337–350.

Garza, R. T., & Borchert, J. E. (1990). Maintaining social identity in a mixed-gender setting: Minority/majority status and cooperative/competitive feedback. Sex Roles, 22(11), 679–691.

Hamilton, W. D. (1964). The genetical evolution of social behaviour. II. Journal of Theoretical Biology, 7(1), 17–52.

Holme, P., & Zhao, J. (2007). Exploring the assortativity-clustering space of a network’s degree sequence. Physical Review E—Statistical, Nonlinear, and Soft Matter Physics, 75(4), 046111.

Hoppitt, W. J. E., & Farine, D. R. (2018). Association indices for quantifying social relationships: how to deal with missing observations of individuals or groups. Animal Behaviour, 136, 227–238.

Kappel, S., Hawkins, P., & Mendl, M. T. (2017). To group or not to group? Good practice for housing male laboratory mice. Animals, 7(12), 88.

Karavanich, C., & Atema, J. (1998). Individual recognition and memory in lobster dominance. Animal Behaviour, 56(6), 1553–1560.

Krause, J., & Ruxton, G. (2010). Important topics in group living. Social Behaviour: Genes, Ecology and Evolution, 203–225.

Macdonald, D. W. (1983). The ecology of carnivore social behaviour. Nature, 301(5899), 379–384.

Newman, M. E. J. (2004). Analysis of weighted networks. Physical Review E—Statistical, Nonlinear, and Soft Matter Physics, 70(5), 056131.

Newman, M. E. J. (2006). Modularity and community structure in networks. www.pnas.orgcgidoi10.1073pnas.0601602103

Oliveira, F. G., Monarca, R. I., Rychlik, L., Mathias, M. da L., & Tapisso, J. T. (2021). Social thermoregulation in Mediterranean greater white-toothed shrews (Crocidura russula). Behavioral Ecology and Sociobiology, 75(10), 147.

Paul, M., & Bhadra, A. (2018). The great Indian joint families of free-ranging dogs. PLoS ONE, 13(5). 10.1371/journal.pone.0197328

Paul, M., Majumder, S. Sen, & Bhadra, A. (2014). Grandmotherly care: A case study in Indian free-ranging dogs. Journal of Ethology, 32(2), 75–82. 10.1007/s10164-014-0396-2

Range, F., & Marshall-Pescini, S. (2022). The socio-ecology of free-ranging dogs. Wolves and Dogs: Between Myth and Science, 83–110.

Sen Majumder, S., Chatterjee, A., & Bhadra, A. (2014). A dog’s day with humans-time activity budget of free-ranging dogs in India (Vol. 106, Number 6).

Shizuka, D., & Farine, D. R. (2016). Measuring the robustness of network community structure using assortativity. Animal Behaviour, 112, 237–246.

Silk, J. B. (2007). The adaptive value of sociality in mammalian groups. Philosophical Transactions of the Royal Society B: Biological Sciences, 362(1480), 539–559. 10.1098/rstb.2006.1994

Stockley, P., & Bro-Jørgensen, J. (2011). Female competition and its evolutionary consequences in mammals. Biological Reviews, 86(2), 341–366.

Sueur, C., Petit, O., De Marco, A., Jacobs, A. T., Watanabe, K., & Thierry, B. (2011). A comparative network analysis of social style in macaques. Animal Behaviour, 82(4), 845–852.

Tibbetts, E. A., Pardo-Sanchez, J., & Weise, C. (2022). The establishment and maintenance of dominance hierarchies. Philosophical Transactions of the Royal Society B, 377(1845), 20200450.

Trivers, R. L. (2017). Parental investment and sexual selection. In Sexual selection and the descent of man (pp. 136–179). Routledge.

Vanak, A. T., Thaker, M., & Gompper, M. E. (2009). Experimental examination of behavioural interactions between free-ranging wild and domestic canids. Behavioral Ecology and Sociobiology, 64(2), 279–287. 10.1007/s00265-009-0845-z

Wey, T., Blumstein, D. T., Shen, W., & Jordán, F. (2008). Social network analysis of animal behaviour: a promising tool for the study of sociality. In Animal Behaviour (Vol. 75, Number 2, pp. 333–344). 10.1016/j.anbehav.2007.06.020

Whitehead, H. (2008a). Analyzing animal societies: quantitative methods for vertebrate social analysis. University of Chicago Press.

Whitehead, H. (2008b). Precision and power in the analysis of social structure using associations. Animal Behaviour, 75(3), 1093–1099.

Whitehead, H. (2009). SOCPROG programs: Analysing animal social structures. Behavioral Ecology and Sociobiology, 63(5), 765–778. 10.1007/s00265-008-0697-y

Willis, C. K. R., & Brigham, R. M. (2007). Social thermoregulation exerts more influence than microclimate on forest roost preferences by a cavity-dwelling bat. Behavioral Ecology and Sociobiology, 62(1), 97–108.

